# Identification of serotonin as a gut regulator of liver hepcidin expression

**DOI:** 10.1101/2021.09.16.460464

**Authors:** Tereza Coman, Marion Falabrègue, Julien Rossignol, Pierre Bonneau, Morgane Djebar, Amélie Bigorgne, Jacques RR Mathieu, Marc Foretz, Léon Kautz, Katel Peoc’h’, Jean-Marie Launay, Luc Maroteaux, Carole Peyssonnaux, Olivier Hermine, Francine Côté

## Abstract

Iron is essential to key biological processes of all living organisms. Proper iron levels must be maintained to meet biological needs and prevent toxicity. Given the central role played by the hormone hepcidin in systemic iron homeostasis, extensive research has sought to identify regulators of its expression. Diverse evidence shows the gut to be an essential sensor and regulator of iron homeostasis, independently of other known hepcidin regulators, including bone marrow signals. Here we identify gut-derived serotonin as a key physiological factor in hepcidin regulation. In response to hypoxia, serotonin synthesized and secreted by enterochromaffin cells can act beyond the gut to repress hepcidin expression in the liver, through a 5-HT_2B_ receptor-dependent pathway. Bone marrow transplant experiments clearly indicate the gut is responsible for hepcidin repression. This regulatory system appears to be conserved in humans: a significant negative correlation exists between hepcidin and serotonin levels in the serum of healthy individuals. Our findings imply hepcidin regulation by serotonin is a physiological process, and modulation of the gut serotonergic system may have broad therapeutic implications.

## Introduction

Abnormally low or high iron levels account for a heavy disease burden worldwide, as they are tied to worse prognoses and higher treatment costs. Thus, iron overload plays a major role in free radical pathology and is linked to tissue damage; heart, liver, or other organ failure; and in some cases, premature death; while iron deficiency, the most common nutritional disorder globally, is associated with anemia as well as adverse pregnancy and birth outcomes (Pasricha et al, 2021; Ganz, 2013; Steinbicker & Muckenthaler, 2013; Kohgo et al, 2008). To prevent and improve the treatment of iron-related disorders, the search for novel regulators of iron homeostasis continues.

Hepcidin, encoded by the *Hamp1* gene and mainly produced by liver hepatocytes, is the central hormone regulating systemic iron homeostasis (Nicolas *et al*, 2002). Highly conserved between species, it decreases intestinal iron absorption and lowers plasma iron concentrations by binding to ferroportin (SLC40A1), the only known iron exporter, on the basolateral surface of duodenal enterocytes (Nemeth *et al*, 2004). Due to the key role played by hepcidin in systemic iron homeostasis, there have been extensive efforts to identify factors regulating its expression (Roth *et al*, 2019; Ramos *et al*, 2011). TFR1 signaling in response to excess iron and IL-6 signaling in the presence of chronic or acute inflammation both stimulate hepcidin production. In contrast, hepcidin is repressed as a result of iron deficiency, hypoxia, and erythroid expansion. Erythroferrone (ERFE/Fam132b)—the long-sought erythroid regulator of hepcidin, released by bone marrow erythroid progenitors in response to acute erythropoietin (EPO) stimulation—acts as a repressor of hepcidin to increase iron availability during the erythroid expansion (Kautz *et al*, 2014).

Besides the erythroid regulator, several pieces of evidence suggest that iron absorption should occur independently of any erythropoietic response in the bone marrow. Tight control of intestinal iron metabolism for maintenance of whole-body iron levels is needed to prevent iron deficiency and toxic iron overload, and is mainly affected by modulation of iron absorption in the proximal gut (primarily in the duodenum). The authors and others have demonstrated that the gut, which has a unique physiological hypoxia profile, regulates the expression of key genes and qualifies as an independent regulator of iron absorption (Shepherd, 1982; Taylor & Colgan, 2007; Mastrogiannaki *et al*, 2009; Shah *et al*, 2009; Pocock & Hobert, 2010). Gut enterochromaffin cells, which express HIF1-α, the canonical regulator of hypoxic responses, have been shown to detect oxygen fluctuations. Moreover, under hypoxic conditions, there is major upregulation of tryptophan hydroxylase 1 (*TPH1*), the gene responsible for synthesis of serotonin (i.e., 5-hydroxytryptamine, or 5-HT), suggesting that HIF1-α may directly drive 5-HT production in cells subject to hypoxia (Haugen *et al*, 2012).

While a study over fifty years ago suggested a role of 5-HT in iron absorption (Thompson, 1965), to this day the connection between iron and 5-HT has remained unclear. 5-HT has been extensively studied as a neurotransmitter, yet gut enterochromaffin cells produce >10 times more 5-HT than the CNS, and gut 5-HT accounts for 90% of peripheral 5-HT (Walther *et al*, 2003; Côté *et al*, 2003). Here we identify gut-derived 5-HT as the factor linking the gut to the liver, repressing the main regulator of iron homeostasis. Specifically, 5-HT produced by enterochromaffin cells in response to physiological hypoxia regulates hepcidin expression in the liver, independently of any erythropoietic activity in the bone marrow.

## Results and Discussion

### Negative correlation between serotonin and hepcidin levels in healthy individuals

We previously showed that mice deficient for gut-derived 5-HT (*Tph1* KO) present a phenotype reminiscent of refractory anemia with ring sideroblasts (RARS) (Malcovati & Cazzola, 2013), a low-risk myelodysplastic subtype characterized by early, substantial iron overload (Sibon *et al*, 2019). We further observed reduced circulating 5-HT in a cohort of low-risk myelodysplastic patients, most of whom had RARS (Sibon *et al*, 2019). Accordingly, our first research aim was to determine whether individual differences in blood 5-HT levels were correlated with hepcidin levels, in a cohort of 89 healthy individuals. Indeed, a significant negative correlation was found (*P* = 0.0041) (**Fig 1A**). Next, the HepG2 cell line—a human hepatoma cell line that frequently serves as an in vitro alternative (Donato *et al*, 2015) to primary human hepatocytes—was used to test whether 5-HT could reduce hepcidin expression. As seen in **Fig 1B**, within HepG2 cells exposed to 5-HT or a 5-HT receptor agonist, there is a significant reduction in the expression of the *HAMP* gene.

**Figure 1.**
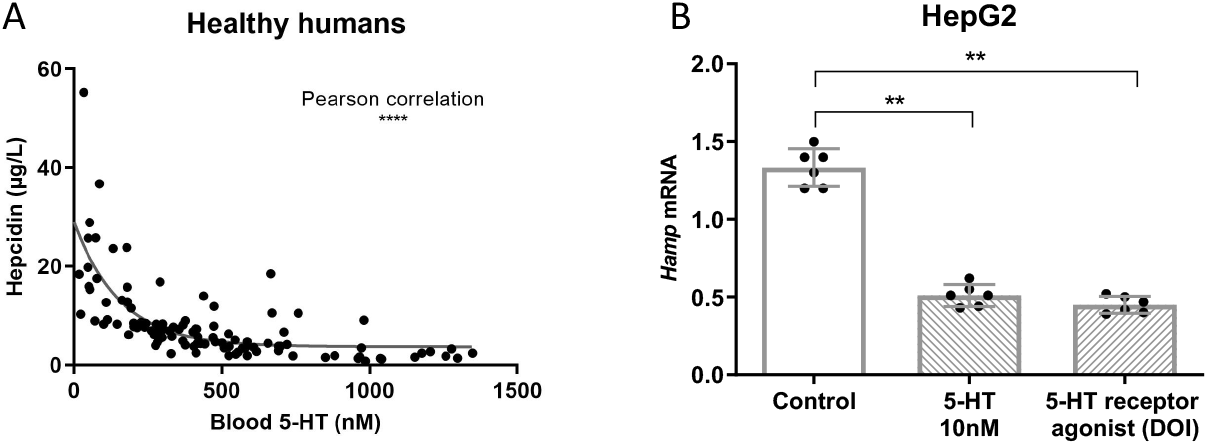
Negative correlation between serotonin and hepcidin levels in humans. A Negative correlation between circulating serotonin (5-HT) and hepcidin in blood samples from 89 healthy humans. B qPCR analysis, normalized against beta-actin, of *Hamp1* expression in HepG2 cell line cultured in vitro with serotonin (5-HT) or 5-HT receptor agonist (DOI). (Data pooled from two independent experiments.) Data information: Individual values presented in addition to means ± standard deviation. Unpaired Mann-Whitney test or Pearson correlations used as appropriate. **: *P* < 0.05; ****: *P* < 0.0001.

### Serotonin represses hepcidin expression through 5-HT_2B_ receptor–dependent pathway

To explore the role played by 5-HT on hepcidin expression, we further analyzed the mouse model deficient for peripheral 5-HT (*Tph1* KO). In *Tph1* KO mice, serum and liver iron (**Fig 2A and B**) and transferrin concentrations (**Fig 2C**) were in the normal range. Ferritin concentration, though significantly higher (**Fig 2D**), was still abnormally low in *Tph1* KO mice compared to what one would expect with RARS. When investigating hepcidin concentrations, we observed a significant increase in steady-state *Hamp1* expression in *Tph1* KO mouse livers, again contrary to what is expected for RARS (**Fig 2E**). Hepcidin increased in these mice despite high plasma EPO concentrations (Amireault *et al*, 2011) and no signs of inflammation (Chabbi-Achengli *et al*, 2016) (**Fig 2F)**.

**Figure 2.**
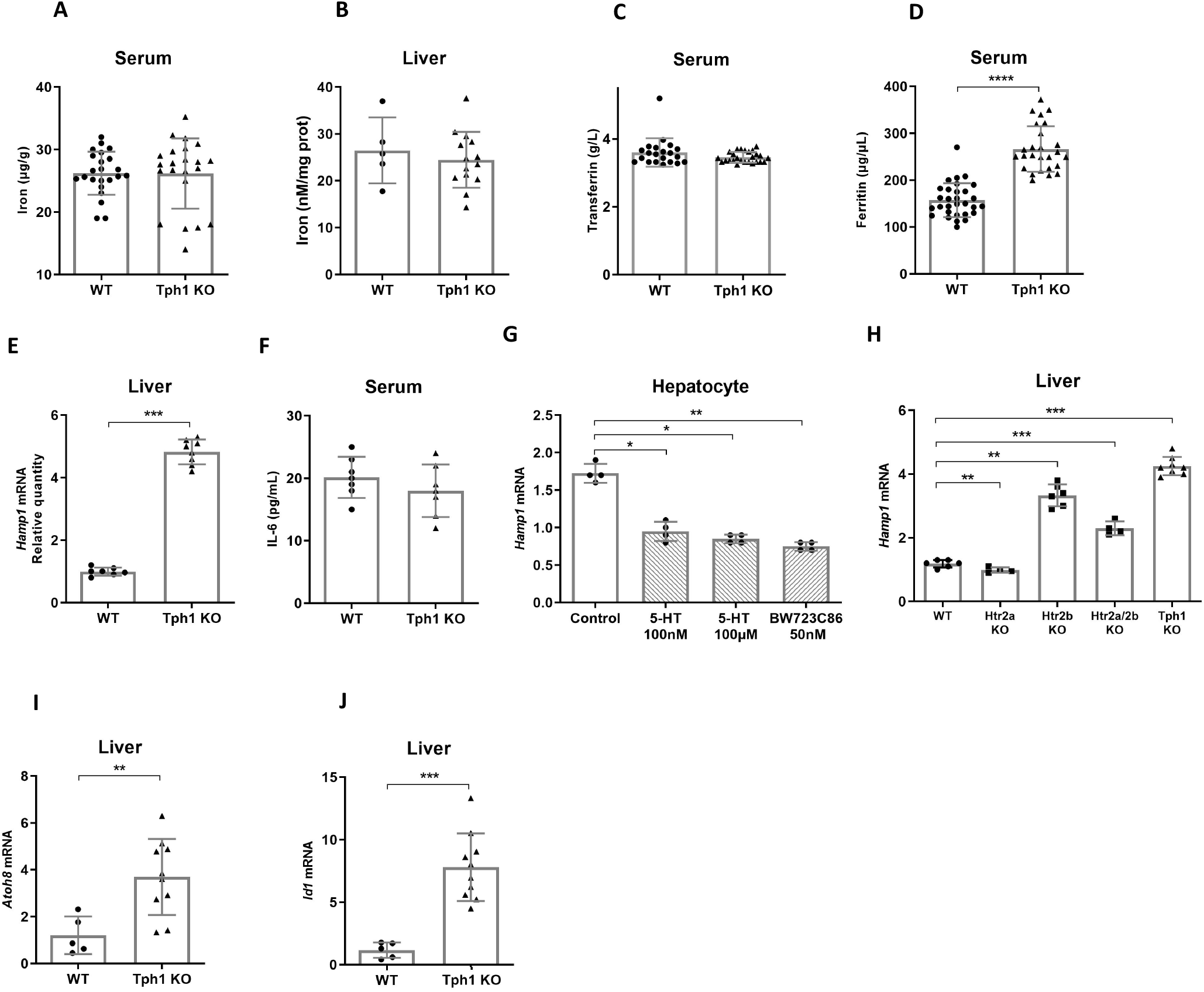
Serotonin represses hepcidin’s expression through a 5-HT_2B_ receptor-dependent pathway. A-F Serum iron (A), total liver iron content (B), serum transferrin (C), serum ferritin (D), total liver *Hamp1* expression (qPCR) (E), and serum IL-6 (F) in wild-type (WT) and *Tph1* knockout (KO) mice. G *Hamp1* expression (qPCR) in murine primary hepatocytes cultured in vitro with serotonin (5-HT) or a 5-HT_2B_ receptor agonist (BW723C86). Data from two independent experiments. H Liver *Hamp1* expression (qPCR) in WT, *Tph1* KO, *Htr2a* KO, *Htr2b* KO, and *Htr2a/2b* KO mice. I, J *Atoh8* (I) and *Id1* (J) expression (qPCR) in livers of WT and *Tph1* KO mice. Data information: The qPCR analyses of *Hamp1* expression were normalized against beta-actin. Individual values presented together with means ± standard deviation. Mann-Whitney tests performed where appropriate. *: *P* < 0.05; **: *P* < 0.005; ***: *P* < 0.0005; ****: *P* < 0.0001.

These human and mouse data led us to hypothesize that 5-HT was a physiological inhibitor of hepcidin. To test this, we first investigated whether 5-HT could directly regulate hepcidin expression in vitro. We and others previously demonstrated the presence of the 5-HT_2B_ receptor (5-HT_2B_R) on hepatocytes (Ebrahimkhani *et al*, 2011). Wild-type (WT) murine primary hepatocyte cell cultures were treated with 5-HT or BW723C86, a 5-HT_2A/2B_ receptor agonist (Diaz *et al*, 2012). *Hamp1* mRNA levels decreased by almost 60% as early as 4 h following either treatment (**Fig 2G)**. To further confirm the involvement of 5-HT_2B_R signaling in hepcidin repression, we measured ex vivo *Hamp1* expression in livers of mice deficient in 5-HT_2A_R, 5-HT_2B_R, and 5-HT_2A/2B_R. Levels of expression in mice lacking 5-HT_2B_R or 5-HT_2A/2B_, as in *Tph1* KO mice, were significantly higher than those in 5-HT_2A_R-deficient mice, which were similar to WT animals (**Fig 2H**).

The mechanism through which 5-HT suppressed *Hamp1* expression was then investigated. It is well established that BMP signaling—involving BMP6, which activates the Smad_1/5/8_ pathway through interaction with its cognate receptor—induces hepcidin production (Meynard *et al*, 2009). We previously reported (Kautz *et al*, 2008) that expression of the genes *Id1* and *Atoh8*, both targets of Smad signaling, constituted a major hepcidin induction pathway. Furthermore, 5-HT inhibition of BMP signaling has previously been demonstrated (Long *et al*, 2006). As illustrated in **Fig 2I and J**, liver basal expression of *Id1* and *Atoh8* was significantly higher in *Tph1* KO mice than in WT mice. This suggests abnormal activation of the BMP signaling pathway in the absence of 5-HT, indicating that a 5-HT signal via the 5-HT_2B_ receptor controls hepcidin expression by modulating BMP/Smad-induced *Hamp1* activation.

### Gut-derived serotonin is key signal in hepcidin regulation in vivo, independently of signals from bone marrow

Liver hepcidin is a central regulator of iron homeostasis, and ERFE/Fam132b has been identified as the erythropoiesis-specific factor affecting hepcidin expression (Kautz *et al*, 2014). Yet tight control of gut iron metabolism, determined by local parameters (e.g., hypoxia), is essential to prevent iron deficiency and toxic iron overload. As (i) enterochromaffin cells are the source of 90% of peripheral 5-HT (Walther *et al*, 2003; Côté *et al*, 2003; Gershon & Margolis, 2021), (ii) hypoxia-responsive elements (HREs) are present in the *Tph1* promoter region (Haugen *et al*, 2012; Sibon *et al*, 2019), and (iii) 5-HT production and secretion are known to be upregulated through a HIF1-α–sensitive mechanism (Haugen *et al*, 2012), we hypothesized that *Tph1* and 5-HT synthesis induced by physiological, or experimental, hypoxia in enterochromaffin cells could constitute a gut-liver axis for *Hamp1* suppression in hepatocytes. Using a mouse model of hypoxia, we observed higher levels of *Tph1* mRNA (**Fig 3A**) and 5-HT (**Fig 3B**) in the duodenum, but not the colon, with a peak at 6 h. Additionally, 5-HT was not metabolized at its production site, as revealed by low concentrations of its 5-HIAA metabolite (**Fig 3C**), and the concentration of 5-HT in the portal vein increased after exposure to hypoxic conditions (*P* = 0.0886) (**Fig 3D**). The portal vein directly connects the gut and the liver, providing abundant amounts of 5-HT for direct and rapid signaling to hepatocytes. Moreover, other authors have shown that portal blood 5-HT concentrations reach a level high enough to exert endocrine effects on the liver (Choi *et al*, 2018; Young *et al*, 2018).

**Figure 3.**
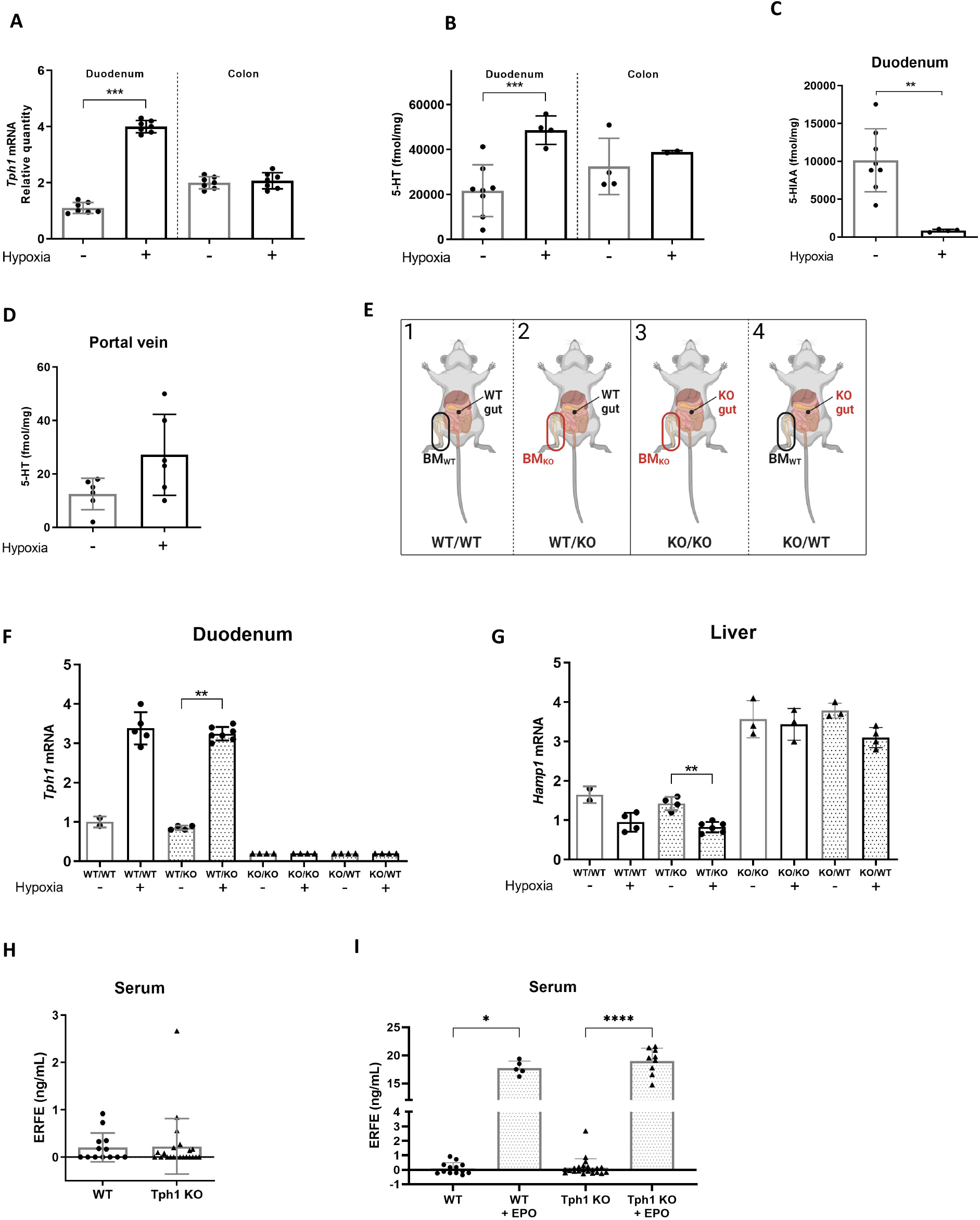
In vivo, gut-derived serotonin essential to regulation of hepcidin expression, independently of any signals from bone marrow. A Duodenal and colonic *Tph1* expression (qPCR) in wild-type (WT) mice exposed to normoxic or hypoxic conditions for 6 h. B-D Duodenal and colonic serotonin (5-HT) (B), duodenal 5-HIAA (C), and portal vein 5-HT (D) levels in WT mice exposed to normoxic or hypoxic conditions for 6 h. E-G Bone marrow (BM) transplant experiments (E) showing relative expression of *Tph1* in duodenum (F) and of *Hamp1* in liver (G) of BM-transplant mice exposed to normoxic or hypoxic conditions for 6 h. WT/WT: WT gut, WT BM; WT/KO: WT gut, *Tph1* KO BM; KO/KO: *Tph1* KO gut / *Tph1* KO BM; KO/WT: *Tph1* KO gut, WT BM. H, I Serum erythroferrone (ERFE) in WT and *Tph1* KO mice at baseline (H) and after EPO injection (I). Data information: The qPCR analyses of *Hamp1* and *Tph1* expression were normalized against beta-actin. Individual values presented together with means ± standard deviation. Mann-Whitney tests performed where appropriate. *: *P* < 0.05; **: *P* < 0.005; ***: *P* < 0.0005.

To firmly establish that peripheral 5-HT regulates hepcidin expression, we next performed bone marrow transplantation experiments as detailed in **Fig 3E**. Briefly, mice had (i) normal gut and bone marrow 5-HT levels (WT/WT), (ii) normal gut 5-HT levels and 5-HT–deficient bone marrow (WT/KO), (iii) 5-HT–deficient guts and bone marrow (KO/KO); or (iv) 5-HT–deficient guts and normal 5-HT bone marrow levels (KO/WT). We measured duodenal *Tph1* mRNA expression at 3, 6, 12 and 24 h of hypoxia, observing peak expression at 6 h. As expected, following hypoxia, a significant increase in *Tph1* mRNA was observed in mice with WT guts, regardless of whether they received WT or *Tph1* KO bone marrow transplants (**Fig 3F**). Furthermore, a significant decrease in *Hamp1* expression was observed in the liver of WT/KO mice, offering further evidence of the role played by gut-derived 5-HT On the other hand, as expected, in KO/KO and KO/WT mice there was no increase in gut *Tph1* mRNA (**Fig 2F**, right) nor decrease in *Hamp1* expression (**Fig 3G,** right).

Importantly, we previously demonstrated that 5-HT was produced in bone marrow proerythroblasts following EPO stimulation (Sibon *et al*, 2019). However, after such stimulation, no mRNA for *Tph1* or 5-HT synthesis was observed within the gut of WT animals. Hence, 5-HT synthesis observed in the gut is a response to hypoxia alone. To further rule out the possibility that bone marrow–derived signals are responsible for hepcidin repression in the liver, we first demonstrated that ERFE is present in the bone marrow cells of *Tph1* KO mice (**Fig 3H**). Secondly, as **Fig 3I** clearly shows, following EPO stimulation bone marrow cells from *Tph1* KO animals could synthesize ERFE at levels identical to those observed in WT mice. Therefore, the inability of KO/WT mice to suppress liver *Hamp1* expression was due to the lack of gut-derived 5-HT—and not of ERFE, which they could still produce.

Overall, the genetic and biochemical findings presented here clearly demonstrate that the observed decrease in *Hamp1* expression under hypoxia is due to gut-derived 5-HT, and appears to occur at 6 h, preceding the erythropoietic regulation of hepcidin, which occurs at 9 to 24 hours according to Kautz et *al* (2014). While ERFE levels have been shown to increase in response to EPO stimulation in acute anemia, mice lacking ERFE have normal hepcidin concentrations at baseline (Coffey *et al*, 2018); this supports the view that ERFE is a stress-induced regulator, rather than a steady-state one. In contrast, as 5-HT-defìcient mice display high hepcidin levels, 5-HT appears to be a physiological regulator of liver hepcidin. Still, it is worth noting that these findings do not exclude the possibility of a time-dependent convergence or complementary mechanisms of bone marrow and gut signals.

Since the discovery of a second gene responsible for 5-HT synthesis (Walther *et al*, 2003; Côté *et al*, 2003), several groups of researchers have demonstrated key physiological roles of nonneuronal 5-HT in several important processes, including hematopoiesis, metabolic homeostasis, and bone metabolism (Gershon, 2013; Mawe & Hoffman, 2013; Amireault *et al*, 2013; Spohn & Mawe, 2017). Our findings are of value to the medical community as they argue for the importance of patient monitoring to prevent and detect potential side effects on iron metabolism exerted by currently prescribed treatments that are known to directly modify 5-HT levels (Hörsch *et al*, 2022; Knupp *et al*, 2022; Sharawat *et al*, 2021).

## Materials and Methods

### Liver iron measurement

Liver iron content was measured by a colorimetric ferrozine assay modified from Riemer *et al* (2004). Tissues were mechanically lysed in 50mM NaOH. The lysate was heated at 60 °C for 2 h in a buffer with equal volumes of 1.4 M HCl and 4.5% (w/v) KMnO4. After cooling samples, iron-detection reagent (6.5 mM ferrozine, 6.5 mM neocuproine, 2.5 M ammonium acetate, and 1 M ascorbic acid dissolved in water) was added to each. After 30 min, absorbance was measured at 560 nm for each sample in a microplate reader (Tecan). Measured intracellular iron concentration was normalized against protein content for each tissue as determined by Bradford assay.

### Mouse ERFE immunoassay

Human recombinant monoclonal antibodies to mouse ERFE were produced by Bio-Rad using HuCAL technology. A high-binding 96-well plate (Corning) was coated overnight at 4 °C with 100 μL/well of 2 μg/mL capture antibody diluted in 50 mM sodium carbonate buffer having a pH of 9.6. The plate was washed (tris-buffered saline, 0.5% Tween 20) and blocked for 1 h with 300 μL/well blocking buffer (phosphate-buffered saline [PBS], 0.2% Na casein, 0.05% Tween 20, 0.1 M NaCl) at room temperature. Recombinant mouse ERFE standard was serially diluted to 10, 5, 2.5, 1.25 and 0.625 ng/mL. Serum samples and standards diluted in PBS and 5% bovine serum albumin (BSA) were added to the wells and incubated for 1 h at room temperature. The plate was then washed and incubated for 1 h with 100 μL/well of biotinylated detection antibody at 0.5 μg/mL in PBS and 5% BSA. After washing, the plate was incubated for 45 min with 100 μL/well of 1:5000 NeutrAvidin–horseradish peroxidase conjugate (Pierce) in PBS and 5% BSA. Next, the plate was developed with 100 mL/well of supersensitive 3,3’,5,5’-tetramethylbenzidine (TMB) substrate (Sigma-Aldrich) in the dark at room temperature. The reaction was stopped by adding 100 mL of 0.5 M sulfuric acid, and absorbance was measured at 450 nm.

### Cell cultures

Primary hepatocytes were isolated from adult wild-type (WT) mice using a collagenase method (Foretz *et al*, 2010). Hepatocytes were seeded in 6-well plates at a density of 300,000 cells per well and cultured under standard conditions (5% CO_2_, 37 °C) in M199 medium containing 2% Ultroser G, for 4 h. After cell attachment, the medium was replaced with fresh M199 medium supplemented with 10% calf serum (Invitrogen).

### RT qPCR

Total RNA was extracted from total duodenum or colon using a RNeasy Kit (QIAGEN). Reverse transcription was performed using iScript Reverse Transcription Supermix for RT-qPCR (Bio-Rad). Real-time PCR was carried out using a StepOne PCR System (Applied Biosystems) with oligos from TaqMan (Life Technologies). Three biological replicates were used for each condition. Data were analyzed with StepOnePlus RT PCR software (v. 2.1) and Microsoft Excel. Beta–actin transcript levels were used for normalization (delta Ct method).

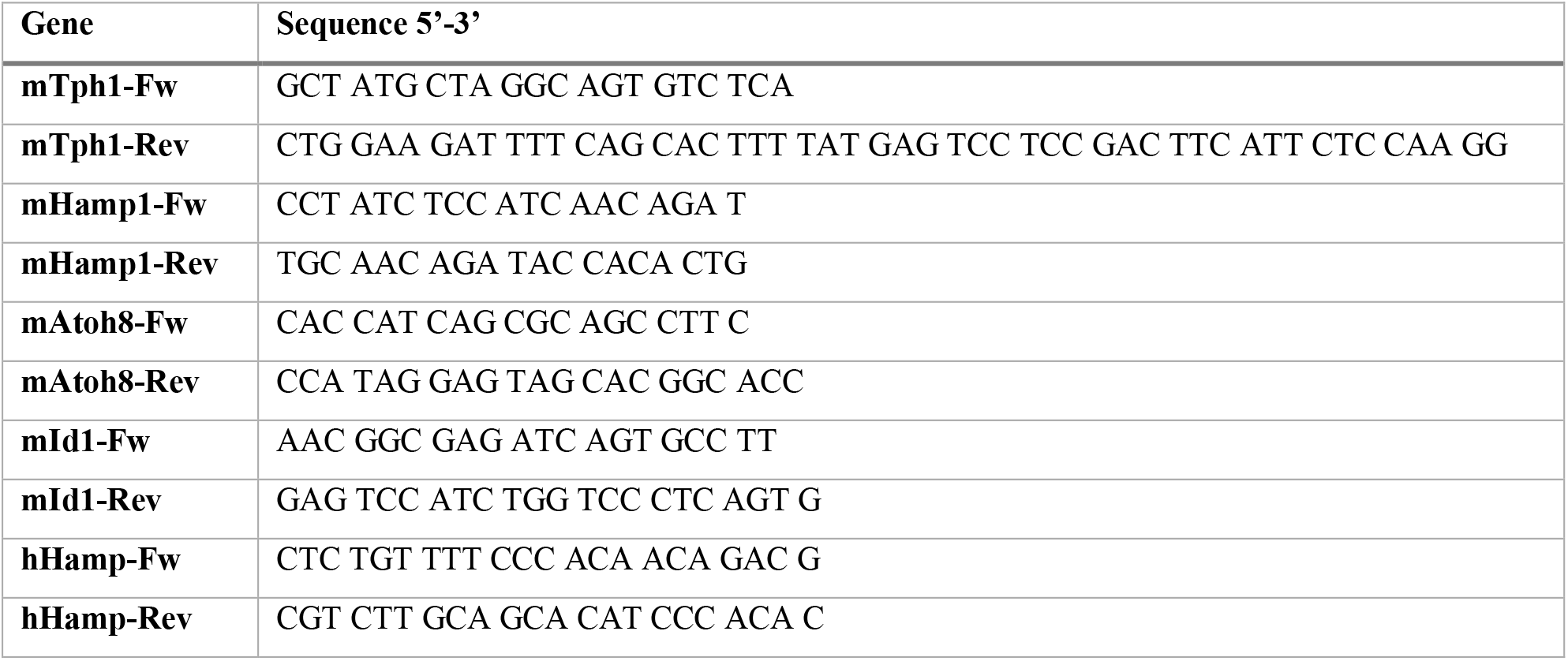

### Animal procedures

*Tph1* KO mice were generated as described by Côté *et al* (2003). *Tph1* KO and WT animals were derived from pure C57BL/6/J (H2b) genetic backgrounds. For some experiments, C57BL/6/J (H2b) mice were purchased from Janvier Labs. *Htr2a* KO, *Htr2b* KO, *Htr2a/2b* KO, and control mice (SVJ129 background) were kindly provided by L. Maroteaux. Male animals were used for each experiment. All mice were bred and housed within a pathogen– free facility, in microisolator cages, and used at 8 to 12 weeks of age. For bone marrow transfer experiments, *Tph1* KO and WT recipients were administered 600 cGy of total-body X-ray irradiation on day −1 as described before (D’Aveni *et al*, 2015). This was followed by intravenous infusion of *Tph1* KO or WT donor bone marrow (BM) cells into the caudal veins of recipient mice, resulting in the following groups of mice: WT/WT (WT gut, WT BM), WT/KO (WT gut, *Tph1* KO BM), KO/KO (*Tph1* KO gut, *Tph1* KO BM), and KO/WT (*Tph1* KO gut, WT BM). Experiments were performed 2 months after the bone marrow transplant procedure.

For hypoxia experiments, WT and *Tph1* KO mice were transferred into a hypoxic chamber at 10% O_2_ (INVIVO_2_ 500, Ruskinn). The treatment lasted between 3 and 24 h. Mice had access to water and food until sacrifice.

Animal experiments were performed according to the recommendations of the ethical review board (Institut Imagine, License A75-15-34).

### Human data

The 89 healthy controls without any hematological disease were recruited from the orthopedic clinic of Lariboisière University Hospital for 5-HT and hepcidin measurements. The study was approved by the local ethics committee and patients provided written informed consent. Fasting blood samples were collected into vacuum tubes containing 109 mM sodium citrate (Greiner Bio-One) with a 9:1 blood to anticoagulant ratio. After removing 0.5 mL of whole blood for 5-HT measurement, the remainder was centrifuged at 3,000 x g for 15 minutes to obtain platelet-poor plasma. The time between blood sampling and plasma isolation was less than 1 hour. Aliquots of whole blood, and plasma were kept frozen (−80°C) until 5-HT and hepcidin measurements. Whole blood 5-HT was measured by high-pressure liquid chromatography coupled to fluorimetric detection (Kema *et al*, 1993) and plasma hepcidin by LC-MS/MS (Lefebvre *et al*, 2015).

Characteristics of the control cohort: age (median, IQR) 38 (29-49); sex M, n (%) 42(47%); risk factors (n, %) diabetes 1(1.1%), arterial hypertension 0(0%), hyperlipidemia 2(2.2%), active smoking 3(3.4%), migraine 1(1.1%), depression 0(0%), history of cardiovascular disease 0(0%), familial history of cardiovascular disease 3(3.4%), history of autoimmune pathology 1(1.1%), combined 2 or >2 risk factors 2(2.2%), none 80(90%).

## Data availability

The datasets produced or analyzed during this study are available from the corresponding author upon reasonable request.

## Acknowledgments

The authors would like to thank Sophie Vaulont for her scientific advice and manuscript review, Aurélia Dujardin and the LEAT Animal Facility for care and breeding of the mice, and Jason Miller for manuscript editing. This research was supported in part by grants from Laboratory of Excellence GR-Ex (ANR-11-LABX-0051), which is funded through the Investissements d’Avenir program of the French National Research Agency (ANR-11-IDEX-0005-02). In addition, TC, MFa, and JR received PhD fellowships from Institut Imagine, Laboratory of Excellence GR-Ex, and the French Ministry of Higher Education and Research, respectively.

## Author contributions

TC, MFa, JR, and FC: conceptualization and formal analysis; TC, MFa, JR, PB, MD, AB, KP, JML, LK, and FC: investigation; JRRM, MFo, and LM: technical support and murine models; FC and OH: funding acquisition; FC: project administration; FC, CP, and OH: supervision; TC, MFa, CP, and FC: writing original draft.

